# NM-RARe: A FRAMEWORK FOR REGIONAL ARTHROPOD CONSERVATION ASSESSMENT: INSIGHTS FROM NEW MEXICO

**DOI:** 10.64898/2026.09.16.752178

**Authors:** Simon M. Doneski, Anna Walker, Quinlyn Baine, Richard Norwood, Thomas P. Bulger, Jade McLaughlin, Matthew L. Forister, Kelly B. Miller, Rayo McCollough, Dewey Devivi, David Lightfoot, Esteban Muldavin

## Abstract

Arthropods represent the most species-rich animal group on the planet, yet they remain largely overlooked in formal conservation planning. New Mexico is a US state with exceptional biodiversity, including numerous endemic, rare, and threatened arthropod species. Recent attention to global insect declines has highlighted the need to document possible threats to and declines of New Mexico insects and other arthropods. All terrestrial arthropods in New Mexico lacked state-level regulatory protection until 2025, and baseline data on the taxonomy, ecology, geographic distribution and conservation needs for most New Mexico arthropod species remained absent. To address this gap, we developed the New Mexico Rare Arthropod Resource (NM-RARe), a publicly accessible regional database for arthropod species and subspecies of conservation concern in New Mexico. Drawing from multiple vetted sources, we compiled an initial working list of 7,018 taxa and conducted 261 preliminary IUCN Red List assessments and 185 NatureServe state rank assessments, covering 285 total taxa. Of those assessed by IUCN Red List standards, 52 were subspecies and 209 were species level assessments, 165 of which met the criteria for inclusion and were incorporated into the NM-RARe database (https://nmrare.org/). Ninety-three of the pollinating insect species from NM-RARe were then formally incorporated into New Mexico’s 2025 State Wildlife Action Plan (SWAP), representing a significant conservation milestone. Beyond its immediate regional impact, NM-RARe presents a standardized, scalable methodology adaptable to any region or taxonomic group, offering a replicable framework for arthropod conservation assessment worldwide.

## Introduction

Concern over global declines in insect populations has grown substantially in recent years, with multiple studies documenting reductions in abundance, diversity, and biomass across a range of taxa and regions (Wagner et al. 2021; Gossner et al. 2023). However, the magnitude and universality of these declines remain contested, as data coverage is geographically and taxonomically uneven (Saunders et al. 2020). Comprehensive, regionally focused assessments are therefore essential to fill these knowledge gaps, evaluate the validity of global “insect apocalypse” claims, and provide land managers and conservation practitioners with reliable information for evidence-based decision making (Duffus et al. 2023).

Conservation of arthropods and assessments like these are particularly critical in ecologically diverse regions such as New Mexico. The state encompasses a remarkable range of habitats and harbors exceptional biodiversity, including numerous endemic species (Danks 1994; Stein 2002). A recent analysis of NatureServe data for New Mexico lists 107 endemic species in the state and many more endemic subspecies (NatureServe 2024). However, NatureServe’s data is not comprehensive, and the real number is likely higher. Despite such high diversity and endemism in the state, formal protection and conservation efforts for insects and arthropods in New Mexico have historically lagged behind those in neighboring southwestern states (Bossart and Carlton 2002).

This historical pattern of neglect for invertebrate conservation in the state has left major gaps in our knowledge and resources needed to conserve arthropods. There is a pervasive lack of basic data on the distribution, life history, and ecology of most arthropod species in the state, particularly narrow range endemics (Montgomery et al. 2020; van der Niet 2021). Because conservation cannot proceed without foundational natural history knowledge, identifying and documenting these species is of critical importance (Start and Gilbert 2019).

To address this knowledge gap, we developed the New Mexico Rare Arthropods Resource ((NM-RARe) found online at https://nmrare.org/). One goal of the project was to conduct an initial assessment of New Mexico’s arthropod fauna to identify species and subspecies that are rare, narrowly endemic, or potentially at risk. A second objective was to develop a publicly accessible, comprehensive database of species of conservation concern to serve as a resource for researchers, land managers, and the public. As part of this, a third objective was to formally assess arthropod species for the IUCN Red List. The IUCN Red List is the globally recognized standard for estimating extinction risk (Rodrigues et al. 2006). In addition, we conducted Natural Heritage New Mexico (NHNM) state status assessments based on population numbers, threats, trends based the NatureServe methodology to feed into their North American database (NatureServe 2026b). To our knowledge, New Mexico is the first state to undertake a project of this kind focused on arthropod information gathering for conservation. It integrates a variety of criteria and assessment methods to develop a more objective definition of rarity and threat of extinction to support state and regional conservation efforts as well as adding to the global perspective for their protection.

This project aimed to fill some of those gaps and builds on a global history of International Union for Conservation of Nature (IUCN) Red Listing initiatives such as the European Red List “Pulse” project which is an EU funded initiative led by the IUCN and over 40 expert institutions to assess 10,000 European species (IUCN 2026). In North America, targeted efforts like the Xerces Society’s *State of the Fireflies* and *State of the Bees* have attempted to Red List assess all species within specific taxonomic groups across the United States (Fallon et al. 2021; Xerces Society 2026). Whereas national and regional Red List initiatives have been undertaken in many places across the world, such efforts have been slow to gain traction in North America. In 1983 the IUCN Invertebrate Red Data Book provided some of the earliest formal assessments of North American arthropods, but it included few New Mexico species and covered only a small fraction of the continent’s arthropod diversity (Wells et al. 1983).

In New Mexico, the only prior attempt to compile such a list was for the 2006 State Wildlife Action Plan, when experts assembled a small list of arthropods based on expert opinion (NMDGF 2006). This listing carried no regulatory status or funding and was effectively symbolic, and it was dropped entirely from the 2016 State Wildlife Action Plan, which retained only aquatic crustaceans and mollusks (NMDGF 2016). The New Mexico Rare Arthropods Resource (NM-RARe) described here builds on the foundation of the 2006 SWAP and other regional assessment projects, advancing their legacy by integrating a range of criteria and assessment methods into a more objective, data-driven definition of rarity and extinction risk. This approach was validated in 2025, when NM-RARe’s assessments supplied the evidence base needed to reinstate arthropods on the SWAP, resulting in the inclusion of 93 insect species on the state’s Species of Greatest Conservation Need list. The first insect species to ever receive the designation in New Mexico.

## MATERIALS AND METHODS

### Compilation of Candidate Species

To initiate development of a database of potentially rare arthropods in New Mexico, a master list of arthropods recorded in the state was compiled from five sources: the original New Mexico 2006 CWCS list (NMDGF 2006), the Biota Information System of New Mexico (BISON-M) (BISON-M 2026), NatureServe Explorer (NatureServe 2024), Cary and Toliver’s (2024) online guide to the butterflies of New Mexico, and the North American Bee Distribution Tool (USFWS 2024). This compilation yielded an initial working list of 7,018 taxa. This is certainly not a comprehensive list, as GBIF.org (2022) includes records for 10,568 arthropod species in New Mexico. However, using the list compiled from the pre-vetted named sources limited the need to verify records across all taxa.

Conservation status ranks were then compiled from NatureServe (2024), the IUCN Red List of Threatened Species (2025), the U.S. Forest Service Regional Forester’s Sensitive Species List (USFS 2013), the Bureau of Land Management Sensitive Species List (BLM 2018), the New Mexico Department of Wildlife’s Species of Greatest Conservation Need List (NMDW 2025), and the U.S. Fish and Wildlife Service Category 2 and Candidate Species lists (USFWS 2026). Of the 7,018 taxa recorded on our initial list, only 246 had a Red List assessment of some kind, and 45 were old assessments, per IUCN guidelines species should ideally be reassessed at least every ten years. These old assessments were excluded this left only 2.86% (201) species from our master list as assessed on the IUCN Red List. Of these, 192 were assessed as Least Concern, one as Endangered, and eight as Data Deficient. Including the old assessments only, 22 arthropod species in New Mexico were assessed as threatened on the IUCN Red List when this project began.

Taxonomists representing a diversity of arthropod groups were then consulted to provide lists of taxa of potential conservation concern in New Mexico, and dozens of experts contributed extensive lists of moths, butterflies, bees, spiders, beetles, aquatic macroinvertebrates, and grasshoppers. Due to resource limitations, a comprehensive assessment of all taxa was not possible at the time. Certain groups were prioritized for inclusion into NM-RARe in this initial effort due to funders’ priorities (i.e. pollinators), available data, and available expertise. For example, butterfly species identified as potentially declining in recent analyses (Forister et al. 2021, 2023; Edwards et al. 2025) were prioritized. Large groups currently underrepresented include most beetle families, true bugs, Chelicerata, and crustaceans. This consultation resulted in a working list of candidate species and subspecies representing a broad taxonomic and ecological range.

### Conservation Assessment

For taxa without recent conservation status assessments or those recommended as rare or potentially threatened by experts, preliminary status assessments were carried out using the categories and criteria of the IUCN Red List and Natural Heritage New Mexico (NHNM) state ranks following the NatureServe methodology.

In the first step of the assessment process, information on taxonomy, distribution, population size and trend, ecology, habitat, behavior, use and trade, threats, and any known or recommended conservation measures was compiled for each taxon from published literature, unpublished reports, and input from taxonomic experts. Occurrence records were obtained from online biodiversity databases and museum collections, including the NM-RARe iNaturalist project, GBIF.org, Ecdysis, Biotics 5, the Museum of Southwestern Biology’s arthropod collection, the scientific literature, and species experts (Ecdysis 2026; GBIF.org 2026; MSB 2026; NatureServe 2026a). Data were screened for erroneous records, which were vetted and removed if unsupported or incorrect. In many cases, records from the published literature and unpublished reports were georeferenced to draft more detailed distribution maps.

We evaluated extinction risk for each species using the IUCN Red List Categories and Criteria: Version 16 as well as with the NatureServe State Rank Criteria using the Conservation Rank Calculator v3.2 (NatureServe 2020, IUCN Standards and Petitions Committee 2024). In the IUCN criteria, each taxa can be assigned to one of the following IUCN Red List categories: Extinct (EX), Extinct in the Wild (EW), Critically Endangered (CR), Endangered (EN), Vulnerable (VU), Near Threatened (NT), Least Concern (LC) or Data Deficient (DD) (Table 1). Our IUCN assessments were preliminary pending IUCN review.

**Table 1.** Comparison of IUCN Red List and NHNM conservation status categories.

| IUCN Categories |  |  | NHNM Categories |  |  |
| --- | --- | --- | --- | --- | --- |
| *Scope: Global* |  |  | *Scope: New Mexico* |  |  |
| Status | Abbr. | Meaning | Status | Abbr. | Meaning |
| Extinct | EX | No wild or captive individuals remain | Presumed Extirpated | SX | Believed extirpated from the state; not located despite intensive searches |
| Extinct in the Wild | EW | Survives only in captivity, cultivation, or as a naturalized population outside its historical range | Possibly Extirpated | SH | Known only from historical occurrences; some possibility of rediscovery |
| Critically Endangered | CR | Very high risk of extinction | Critically Imperiled | S1 | Very high risk of extirpation |
| Endangered | EN | High risk of extinction | Imperiled | S2 | High risk of extirpation |
| Vulnerable | VU | Moderate risk of extinction | Vulnerable | S3 | Moderate risk of extirpation |
| Near Threatened | NT | Low risk of extinction | Apparently Secure | S4 | Low risk of extirpation |
| Least Concern | LC | No risk of extinction | Secure | S5 | No risk of extirpation |
| Data Deficient | DD | Not enough data to rank | Unrankable | SU | Not enough data to rank |

**Table 2.** Criteria for inclusion of taxa on NM-RARe.

| No. | Criterion |
| --- | --- |
| 1 | IUCN ranked globally, in draft or published form, as Critically Endangered (CR), Endangered (EN), Vulnerable (VU), or Data Deficient (DD). |
| 2 | Ranked globally on NatureServe as Critically Imperiled (G1), Imperiled (G2), Vulnerable (G3), or Unrankable (GU). |
| 3 | A subnational rank in New Mexico of Critically Imperiled (S1), Imperiled (S2), or Vulnerable (S3). |
| 4 | Federally Endangered or Threatened Species. |
| 5 | State Endangered or Threatened Species. |
| 6 | New Mexico Department of Wildlife Species of Greatest Conservation Need. |
| 7 | Forest Service Sensitive Species. |
| 8 | Bureau of Land Management Sensitive Species or Watch List Species. |
| 9 | Endemic species to New Mexico. |
| 10 | Lost Species. |
| 11 | Navajo Natural Heritage ranked Critically Imperiled (G1), Imperiled (G2), Vulnerable (G3), or Unrankable (GU). |
| 12 | Consensus for inclusion by the NM-RARE Arthropods Technical Council (generally at the request of an expert). |

**Table 3.** Threat codes checked for 185 NatureServe state rank assessments.

| Threat | Number of Taxa |
| --- | --- |
| Agriculture | 107 |
| Climate Change and Severe Weather | 87 |
| Natural System Modifications | 86 |
| Invasive and Other Problematic Species | 52 |
| Pollution | 38 |
| Residential and Commercial Development | 57 |
| Energy Production and Mining | 20 |
| Human Intrusions and Disturbance | 18 |
| Biological Resource Use | 8 |

With respect to NHNM state ranks they operate very similarly to IUCN ranks were taxa can be assigned to one of the following categories. However, notably our IUCN global ranks look at species or subspecies status at a global scale NHNM state ranks only look at New Mexico and only use threats and occurrence records from the state. As a result, taxa can be assigned to one of the following categories: Presumed extirpated (SX), possibly extirpated (SH), Critically Imperiled (S1), Imperiled (S2), Vulnerable (S3), Apparently Secure (S4), Secure (S5), or Unrankable (SU) (Table 1).

After preliminary Red List assessments and NatureServe state assessments were completed, the taxa were then assessed against the NM-RARe criteria to determine if they could be included on NM-RARe. Taxa were included on NM-RARe when they met any two of the following criteria:

Endemic species were defined as restricted in range to within the political boundaries of New Mexico. Lost Species were defined following the criteria of Forister and Woodward (2026) as those with no detections in the past ten years. Importantly, Lost Species status does not imply extinction or poor conservation outlook but may instead reflect inadequate survey effort. As a result, the status of Lost Species is something of a sliding scale where conservation concern increases with survey effort. Furthermore, several NM-RARe taxa are published and peer reviewed as Lost Species (Doneski and Baine 2026; Doneski and Miller 2026). Besides those 5 taxa currently 43 others assessed by NM-RARE meet the definition of Lost Species but are not published and their status has not been peer reviewed (Appendix. 1). As a result, more concern should be placed on published taxa who have gone through a rigorous review process at the Journal of Lost Species.

### Development of Public Facing Website

Once NM-RARe’s initial round of assessments was completed and a number of taxa were identified as eligible for inclusion, a public-facing website was developed (https://nmrare.org/). The site was initially built on a MySQL database and later migrated to a WordPress platform. Development was coordinated between Natural Heritage New Mexico and the Colorado Natural Heritage Program. Search functionality was added during the initial development phase, allowing users to search by county, taxonomic order, land management agency, conservation status, and common or scientific name. Species maps were pulled from GBIF.org, and species profile pages were written by biologists at the Museum of Southwestern Biology and the New Mexico BioPark Society.

## RESULTS

In total, 261 preliminary IUCN Red List assessments were completed during the NM-RARe process, a number exceeding the total number of Red List assessments that had been completed in New Mexico on arthropods previously (n = 201). These assessments are currently still under revision and have been periodically submitted for publication since 2024, with 25 published so far. This brings the proportion of New Mexico’s described arthropods with a Red List assessment to 6.54% based on our initial list. In addition, 185 NHNM/NatureServe state assessments were completed for New Mexico, all of which have now been published. Combined, NM-RARe produced 446 total assessments covering 285 total taxa (Appendix. 1). Of these, the majority were Lepidoptera, followed by Hymenoptera and Coleoptera (Fig. 1).

**Fig. 1.**
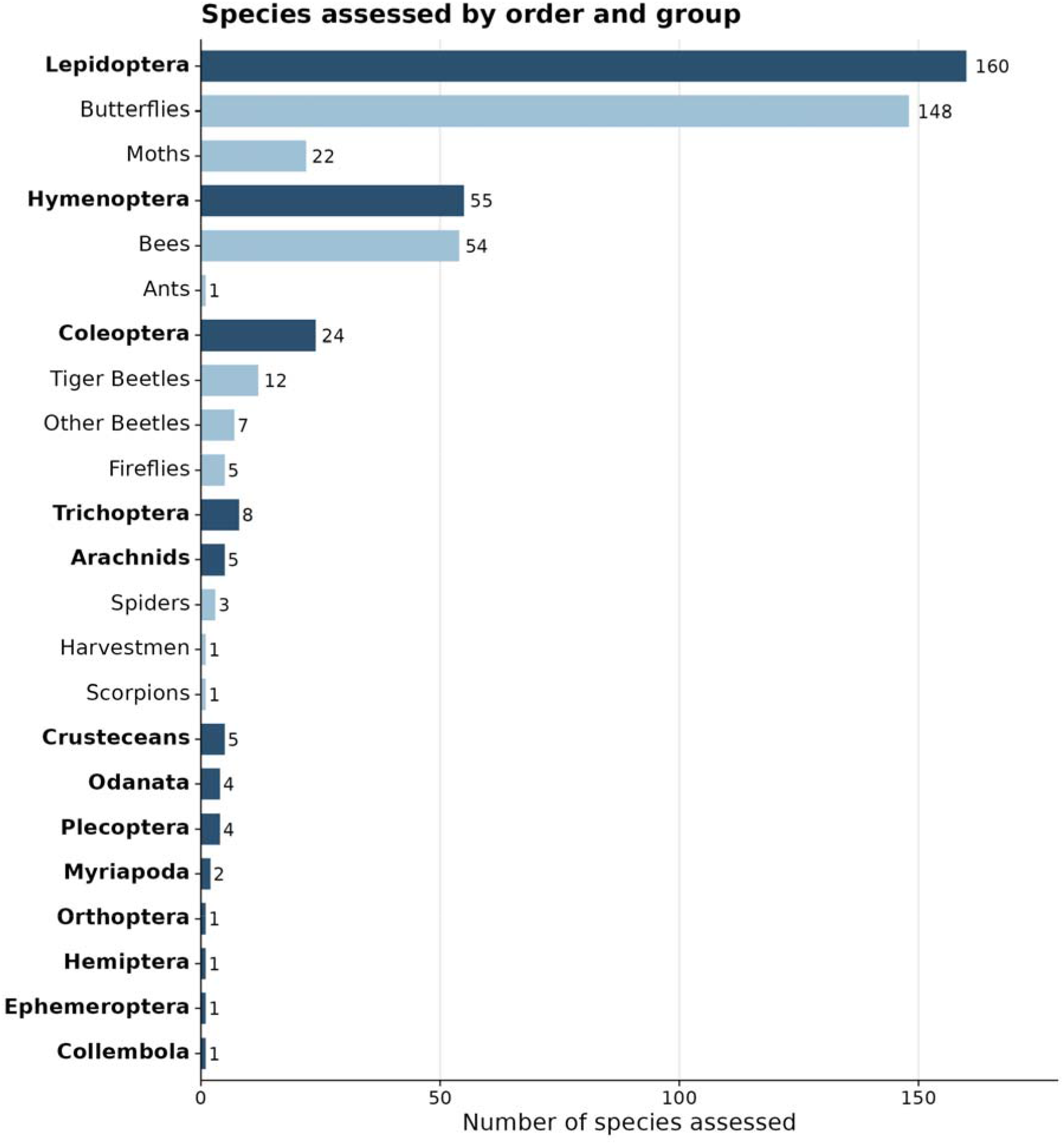
Number of arthropod species assessed under NM-RARe by taxonomic order and group. Orders are shown in descending order of the total number of species and subspecies assessed. Bold and dark blue indicates orders and light blue, and no bold indicates functional groups or suborders.

For the draft IUCN Red List assessments, 56.1% came out as threatened with 36.2% falling as Endangered or Critically Endangered (Fig. 2). The NHNM state ranks were less conservative with 92.8% facing moderate or higher risk of extirpation in the state and 82.4% facing a high or very high risk of extirpation (Fig. 3).

**Fig. 2.**
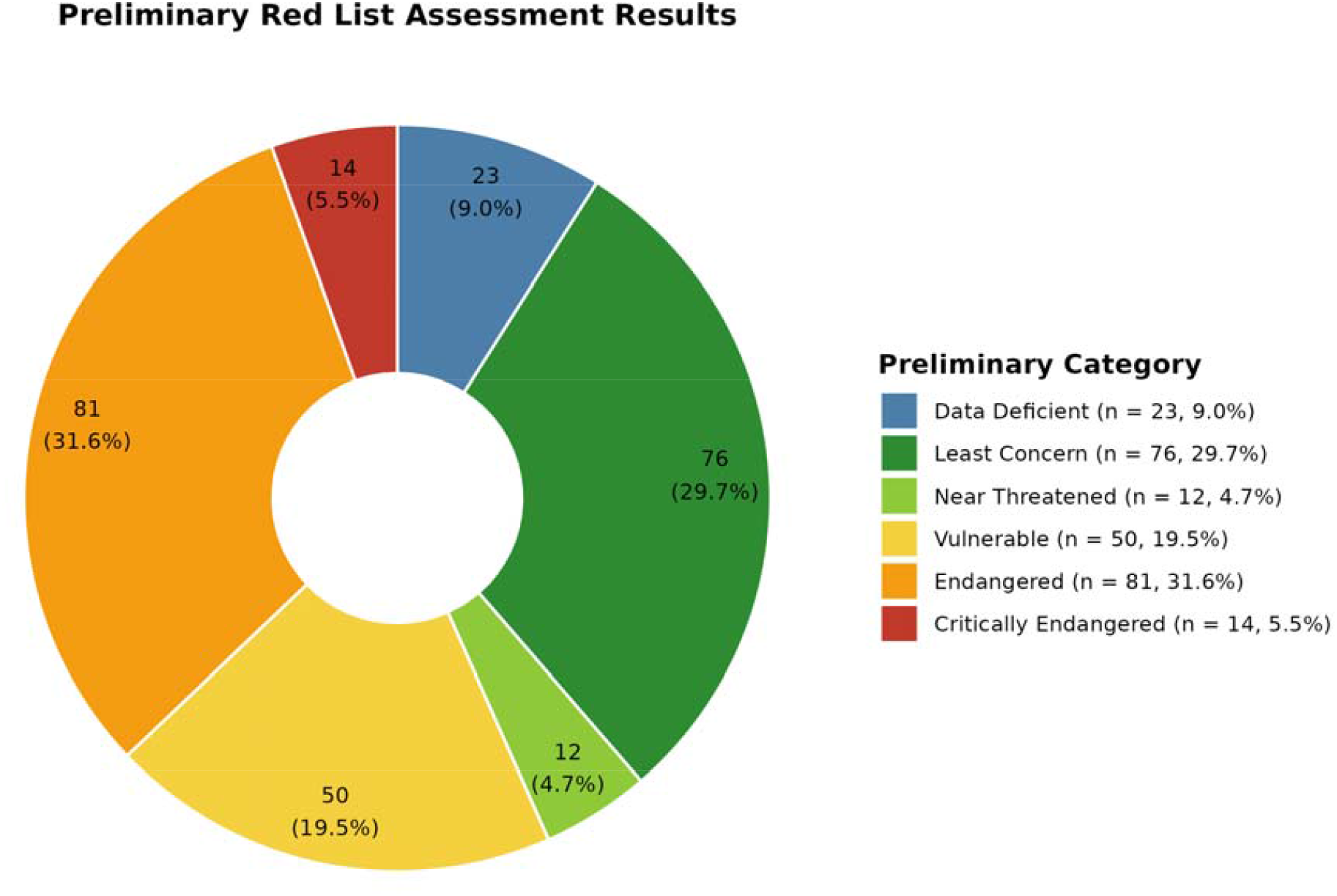
Preliminary IUCN Red List category assigned to each species assessed (n = 256).

**Fig. 3.**
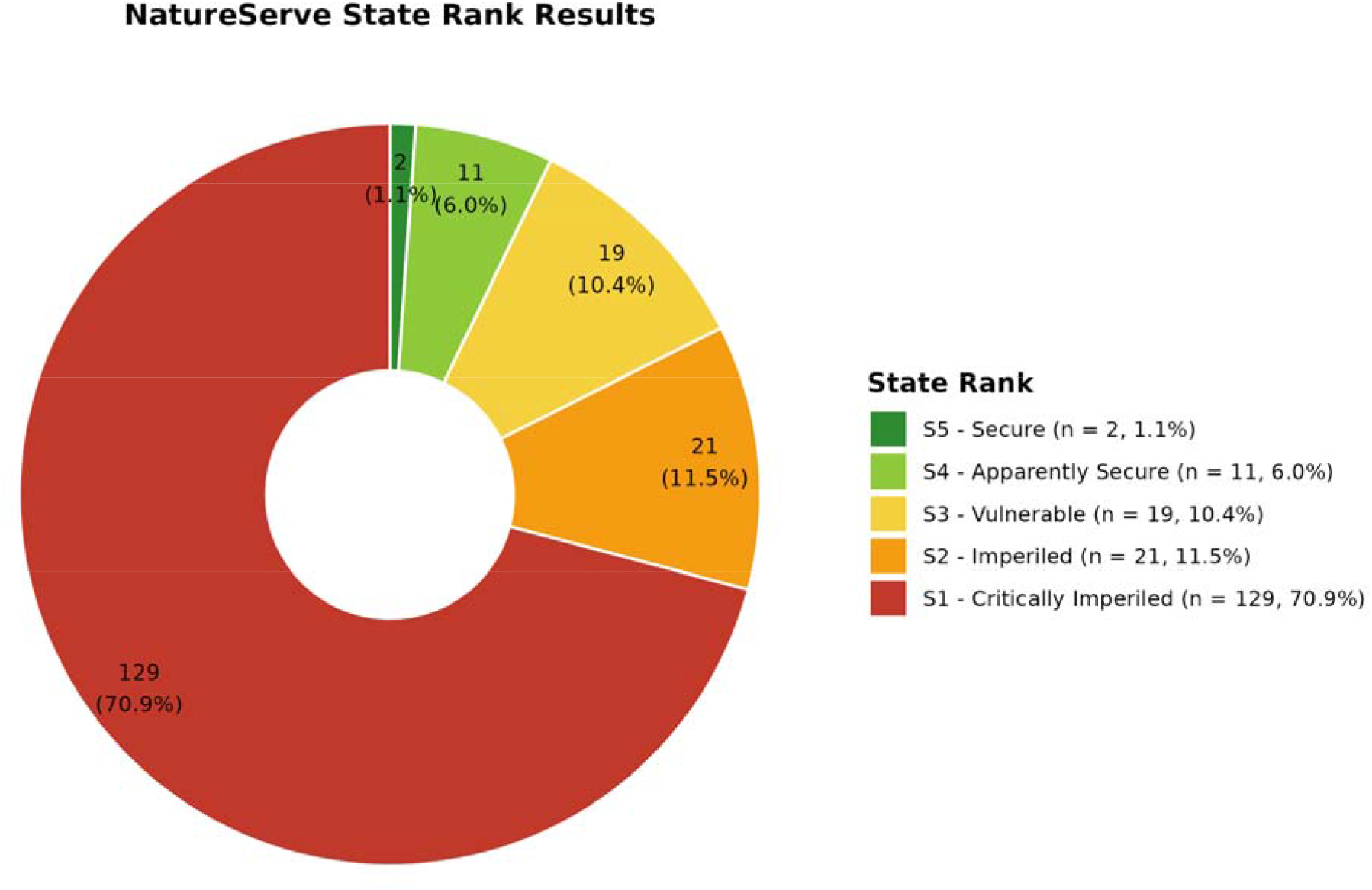
NHHM state ranks assigned to each species (n = 185).

Of the 284 taxa assessed, 165 taxa met the criteria described above for inclusion in the NM-RARe database. This database was designed as a flexible resource that can accommodate revisions as new distributional data, taxonomic updates, and conservation assessments become available. Continued curation and expert review will ensure that the database reflects the most current understanding of arthropod rarity in the region.

Next, we attempted to identify what was causing so many of the arthropod taxa in New Mexico to be threatened with extinction. During NHNM state assessments, threat codes were examined and recorded to do this. For higher level threats the following were used during our NHNM state assessments:

## DISCUSSION

### Assessment Results

Whereas NM-RARe specifically targeted rare species and elevated threat ranks were therefore expected, the proportion of species falling into threatened categories was substantially higher than anticipated, relative to pre-initiative assessments for New Mexico. Over 50% of taxa we assessed using Red List standards and criteria came out in threatened categories (see Fig. 1). This also increased the number of threatened arthropods assessed with the IUCN Red List framework in New Mexico from 23 to 168 taxa, a seven-fold increase. However, much more work on assessments is needed. Our assessments combined with the previously published current assessments cover just 5.8% of the taxa on our initial working list of 7,018. There are also many subspecies which need assessments as well, which are not currently reflected in our list. Furthermore, reassessments for the 22 historical assessments are needed, and as NM-RARe continues, it will need to plan to reassess its own taxa list periodically.

NHNM state ranks following NatureServe guidelines provide a comparable framework but reflect extirpation risk at the state level, which for New Mexico endemic species would mean extinction. Given NM-RARe’s focus on rare taxa within the state, the NHNM state ranks were expected to be and were considerably higher than the corresponding IUCN global ranks (44% more species listed as Vulnerable or higher). We suggest that this reflects a more intensive look at imperilment at the state level that can help guide conservation at local levels which in turn can inform global assessments,

### Threats

Threats to species were categorically identified through the state NatureServe-based ranking process, but these should be interpreted with some caution, given the limited scope of the underlying data. Only 185 taxa or 2% of the original list were evaluated, and a number of these were subspecies rather than full species. Despite the limited taxa coverage, the threat patterns that emerge were informative. Agriculture, which included grazing and pesticide use, was flagged as a prevalent threat across the majority of all arthropod taxa assessed, which is consistent with the extensive agricultural land use and associated chemical application across the state. Climate change also emerged as a leading threat, unsurprising given that many of the assessed taxa are high elevation endemics with narrow thermal tolerances and limited capacity to shift range in response to warming (Rödder et al. 2021). Within the natural systems modifications category, fire suppression and dams or water management were the two threats that recurred most frequently. New Mexico’s arid climate makes its arthropod fauna particularly sensitive to changes in water availability, and even modest reductions in the water table can result in local extirpations, particularly for taxa dependent on riparian or wetland habitat (USFWS 2005). Similarly, decades of fire suppression have altered natural disturbance regimes across the state, contributing to the loss of early successional habitat that many fire-adapted or open-habitat specialist taxa depend on (Kaufmann et al. 1998).

### Taxonomic Bias

A project like NM-RARe represents a long-term commitment rather than a finite undertaking. Arthropods are an extraordinarily diverse group, and assessing species status requires substantial time, effort, and funding, resources that are rarely available in proportion to the scale of arthropod diversity itself. This initial phase of NM-RARe focused on taxa for which the most data were already available, and as a result the current assessment list carries significant taxonomic bias (Fig. 1), with well-studied groups such as butterflies and bees overrepresented relative to their share of the state’s true arthropod diversity. The goal moving forward will be to continue expanding NM-RARe to encompass new taxonomic groups, as well as to conduct additional assessments within groups already included. Achieving anything resembling a truly comprehensive list may be a decades-long process, and in all likelihood the effort will never be fully complete, given the pace at which new taxa are described and reassessed. In the next phase, we plan to further expand NM-RARe to include more in-depth assessment lists for crustaceans, arachnids, Hemiptera, and Coleoptera. However, for many of these groups, significant expertise and data hurdles remain, as well as political hurdles including a shortage of taxonomic specialists and limited baseline distributional data. Overcoming these hurdles may take many years or decades before a comprehensive assessment becomes feasible. There are also significant educational and political hurdles to working on many groups which the public and policy makers have an unfavorable view of such as flies, wasps, and roaches or any insects perceived as agricultural pests. This can make procurement of funding or conservation resources challenging.

### Conservation impact and implications

Effective conservation requires foundational knowledge; we cannot conserve what we do not know. Until recently, New Mexico like many other western states lacked regulatory status on invertebrates. As a result, relatively little attention has been given to them in a conservation context and less foundational knowledge is available than for many other taxa. NM-RARe represents a transformative step forward, providing for the first time a list of arthropods at risk of extinction in the state, supported by rigorous assessments. In 2025, the New Mexico Department of Wildlife gained regulatory authority over invertebrates and, the same year, incorporated the pollinators listed on the NM-RARe website onto their State Wildlife Action Plan as Species of Greatest Conservation Need no insects were given this status before 2025. This designation marked a significant milestone by making these species eligible for agency resources and funding, while elevating their profile in statewide conservation priority-setting. More broadly, this outcome illustrates an important pathway through which taxonomic experts and researchers can translate scientific knowledge into meaningful conservation action.

Since its publication, NM-RARe has expanded to include a Technical Council comprised of taxonomic experts, scientists, and conservationists. This council will oversee the database to ensure it is maintained according to the best available science and provides governance over decisions to add or remove species. In this way, the database remains an objective, science-based resource for agencies and NGOs to incorporate into their conservation planning.

A defining feature of NM-RARe was its use of transparent, objective criteria for assessing rarity and threat status, drawn from multiple publicly available sources. As a result, this standardized framework could be applied anywhere, and it is scalable enough to determine rare species at any geographic or taxonomic range. By integrating multiple indices, the framework also mitigates limitations inherent to any single source. For example, the IUCN Red List can be poorly suited to the needs of insect conservation at a local level. Of its five listing criteria, four require some level of population or trend data, information that is absent for essentially all insects with the exception of a small number of butterfly species (owing to the work of Forister et al. (2021) and Edwards et al. (2025)). This leaves Criterion B “Geographic range in the form of either (extent of occurrence) AND/OR (area of occupancy)” as the only viable pathway to listing for the vast majority of arthropod taxa. Yet even this criterion is difficult to apply consistently to invertebrates, since location-based threat data are often sparse, unknown, or not easily assigned a geographic footprint, as is the case with climate change or drought. A species that is poorly resolved under Red List criteria may nonetheless be identified as rare through one of the other criteria, where threat data with a geographic footprint is not required, as is the case with NatureServe, where a small number of documented occurrences alone can denote a threatened rank However, NatureServe carries its own limitations, in that its assessments are not peer reviewed, and its state ranking process is generally not fully transparent or accessible.Combining these sources therefore reduces the risk of both false negatives, rare species overlooked because they fail to meet any single system’s criteria, and false positives, non-rare species misclassified as threatened due to gaps or biases in one framework alone. This cross-validation increases confidence that genuinely rare species are correctly identified, and ensures that critical designations, such as Lost Species status or those listed as threatened by state or federal agencies, are appropriately prioritized.

NM-RARe is not just a regional assessment project but was also built into a state of the art easy to use website interface. This user-friendly framework makes this data much more accessible to scientists, land managers, and the public than these processes have historically been.

Furthermore, members of the public can submit recommendations for species to be added or removed from the list through this website, ensuring that the website is democratic and constantly evolving. These suggestions are then approved by a council of taxonomic experts and conservationists. Beyond the website, NM-RARe represents a substantial contribution to the scientific and conservation communities through the generation of numerous new IUCN Red List assessments and NatureServe state ranks, adding meaningful data to global and regional biodiversity records, where little previously existed. The data produced by NM-RARe empowers local resource managers, biologists, and scientists with actionable information and concrete steps toward conserving rare arthropod species and subspecies. Whereas the current phase of the project is limited by taxonomic bias and will require decades of continued effort to approach anything resembling comprehensive coverage, the framework established here provides a durable foundation upon which to build. Critically, NM-RARe also presents a standardized, scalable methodology that can be adapted to any geographic or taxonomic scale, and by drawing on multiple complementary data sources, it helps overcome the limitations inherent to any single assessment system. This framework offers a replicable template for how other states, countries, or regions might begin to assess and protect their rare arthropod fauna, an endeavor made ever more urgent in light of reported widespread global arthropod declines.

## Acknowledgements

The authors gratefully acknowledge the Carroll Petrie Foundation and Jennifer Pedneau for their generous support of this project and for making NM-RARe possible. We also extend our sincere thanks to Ginny Seamster at the New Mexico Department of Wildlife, whose contributions were integral to every stage of this work. Furthermore, this work would not have been possible without the efforts and support of taxonomic experts who wrote assessments, reviewed assessments and provided lists and photos for use on NM-RARe. As a result we would like to thank the following: Steven J. Cary, Kevin J. Burls, David L. Wagner, Michael E. Toliver, Charles B. Knisley, Mark S. Romero, Eliza M. Grames, Michael J. Anderson, Candace Fallon, Christopher A. Halsch, Chris C. Nice, Karen Gaines, Rich Hatfield, Cheryl B. Schultz, AndrewD. Warren, Marshal C. Hedin, Chuck Harp, Paul K Masonick, Steve Nanz, Joshua P. Jahner, Vaughn Shirey, Robert M. Pyle, Olivia M. Carril, Jade McLellan, Jim P. Brock, Jerry Jacobi, Monni Böhm, Elliot Gordon, Felix AH Sperling, Doug Yanega, Jeffery Huether, John Gruber, Neil Cox, David L. Pearson, Don B. Thomas, Merrill A. Peterson, Elena S. Tartaglia, Daniel Rubinoff, Erin O. Campbell, Charles Oliver, Brittany Wingert, Oscar Martinez Lopez, Allen Cabrero, Catherine Numa, Leendert-Jan van der Ent, Sophie Ledger, Ale Turkmen, John Shuey, Scott Ellis, Terry Ireland, and Betsy Bainbridge.

